# Evaluating the transmission risk of amyloid beta peptide via ingestion

**DOI:** 10.1101/2023.08.03.551437

**Authors:** Joshua Raine, Nicholas Tolwinski, Jan Gruber, Ajay S. Mathuru

## Abstract

**Background:** Recent reports suggest that amyloid beta (Aβ) peptides can exhibit prion-like pathogenic properties. Transmission of Aβ peptide and the development of associated pathologies after surgeries with contaminated instruments and intravenous or intracerebral inoculations have now been reported across fish, rodents, primates, and humans. This raises a worrying prospect of Aβ peptides also having other characteristics typical of prions, such as evasion of the digestive process. We asked if such transmission of Aβ aggregates via ingestion was possible.

**Methods:** We made use of a transgenic *Drosophila melanogaster* line expressing human Aβ peptide prone to aggregation. Fly larvae were fed to adult zebrafish under two feeding schemes. The first was a short-term, high-intensity scheme over 48 hours to determine transmission and retention in the gut. The second, long-term scheme specifically examined retention and accumulation in the brain. The gut and brain tissues were examined by histology, western blotting, and mass spectrometric analyses.

**Results:** None of the analyses could detect Aβ aggregates in the guts of zebrafish following ingestion, despite being easily detectable in the feed. Additionally, there was no detectable accumulation of Aβ in the brain tissue or development of associated pathologies after prolonged feeding.

**Conclusions:** While human Aβ aggregates do not appear to be readily transmissible by ingestion across species, two prospects remain open. First, this mode of transmission, if occurring, may stay below a detectable threshold and may take much longer to manifest. A second possibility is that the human Aβ peptide may not be able to trigger self-propagation or aggregation in other species. Either possibility requires further investigation, taking into account the possibility of such transmission from agricultural species used in the food industry.

## 1. Background

Hyperphosphorylated tau tangles, amyloid (Aβ) peptide oligomers, and plaques are proposed to play an important role in Alzheimer’s Disease (AD) progression in many AD subjects [1–5]. Although the etiology of the disease is complex and the exact molecular steps leading to AD are unclear [6], reports of cognitive health improvement in clinical trials with monoclonal antibodies targeting Aβ among early symptomatic AD patients have put the spotlight back on amyloid [3,7]. Isoforms of Aβ peptide ranging between 37-43 amino acids are formed from the cleavage of the amyloid precursor protein (APP) at different sites by an assortment of β- and γ-secretases [8]. A long-standing question of interest not only for Aβ peptides but also for other proteins and peptides associated with neurodegeneration such as tau, and α-synuclein, is how pathogenicity spreads during disease progression [9]. Many independent studies support the proposal that Aβ aggregates have prion-like properties such as self-propagation in the disease state [10–14]. An abnormally folded monomer in an alternate confirmation is hypothesized to act as a seed, polymerizing to form paranuclei oligomers, which polymerize further to form fibrils [14–16]. As these insoluble fibrils grow, they are then thought to act as secondary nucleation sites, creating self-replicating amyloid fibrils that aggregate as new plaques with other proteins [11]. Such self-propagation is cited to allow Aβ to act as a pathogen, propagating strains distinct in isoform deposition ratios, and plaque morphology [17].

This self-propagation mechanism becomes problematic when placed in the context of the potential harm to the general public from its transmission, even if it were to occur in only a fraction of AD cases. Alarmingly, reports of transmission of Aβ through medical procedures have been documented in recent years [18–20]. First reported in 2015 for humans, and subsequently confirmed by testing of archived material in 2018, these instances of iatrogenic transmission of Aβ are proposed to have occurred through treatments with contaminated cadaveric pituitary growth hormone [18,19]. The gray matter and pituitary glands of those given the contaminated material exhibited Aβ pathologies typical of early-onset AD while possessing none of the genetic predisposition markers [18,19]. Since this initial report, over 20 individual cases of iatrogenic Aβ transmission resulting in cerebral amyloid angiopathy (CAA) have been reported, categorized by a lower average age of AD onset when compared to sporadic CAA [20].

This pathogenic nature of Aβ has been examined extensively in primates, rodents, and zebrafish [21–25]. Intracerebral inoculation of AD-afflicted human brain homogenates in mouse lemurs, for example, triggered encephalopathy and cognitive decline over two years, with the earliest symptoms occurring after just four months [22,24]. The results were comparable in marmosets for observing cerebral amyloidosis, though it required a longer incubation time of approximately six years [26–28]. Rodent models, on the other hand, showed shorter incubation times of four to six months post-inoculation before significant plaque formation was observed in APP-producing transgenic lines with a predisposition to develop Aβ pathologies [21,25,29,30]. In experiments with zebrafish, there is evidence for Aβ causing plaque-like deposition, cognitive impairment, and truncated lifespans in a time scale of days to weeks following inoculation [31–34].

A notable characteristic of other prions [35–38], which remains untested for Aβ, is the capacity to evade the digestive system and transmit zoonotically by ingestion. The most well-documented examples of such transmission are the transmissible spongiform encephalopathies due to consumption of meat contaminated with bovine spongiform encephalopathy (BSE) causing prions, and the sudden onset of Creutzfeldt–Jakob disease [35,36,39–41]. Critically, recent evidence has highlighted that Aβ injected into the gut tissue of mice is capable of migrating to the brain [42]. In addition to this, misfolded alpha-synuclein, relevant to Parkinson’s disease, exhibits a similar phenomenon when expressed in the gut of *D. melanogaster* [38]. These observations suggest ingestion is a potential route for transmission from the gut to the brain. However, the digestive processes were bypassed in both instances, and thus the question as to whether Aβ can survive this process, and be absorbed unscathed to remain viable as a pathogenic seed is unanswered. Our study was designed to address this question directly.

## 2. Methods

### 2.1. Organism generation and culture

Transgenic *D. melanogaster* expressing either a fluorescent optogenetic human Aβ (Aβ-CRY2-mCherry) or a control fluorescent protein (tdTomato) used for feeding were generated as described previously [23,43]. As noted in previous studies, visible light was sufficient to induce aggregation of Aβ-CRY2-mCherry in *D. melanogaster* larvae [23,43]. Zebrafish (*D. rerio*) were bred and housed as described in Nathan *et al*., 2022 [44] at the ZebraFish Facility (ZFF, Institute of Molecular and Cell Biology, A*STAR) in groups of 20–25 in 3-L tanks under standard facility conditions. Experimental protocols approved by the IACUC committee, Biological Resource Center at A*STAR (IACUC #201571) were followed for experiments with zebrafish.

### 2.2. Feeding schemes and tissue extractions

*D. rerio* were fed dead *D. melanogaster* third instar larvae or extracts (S1 Feeding video). For the short-term intensive feeding scheme to quantify visually, fish were fed individually (with ∼10 larvae). The number of larvae consumed was recorded, and the process was repeated after 24 hours. Fish were euthanized 24 hrs after the second feeding and tissue was harvested for processing or histology. The complete gut tissue, including the intestinal bulb and posterior intestine, was dissected out as described in Gupta and Mullins, 2010 [45], placed in 300 μl RIPA buffer, and homogenized on ice for 1 minute to lyse the cells. The lysed extract was centrifuged at 14000 x G, 4°C, for 10 minutes. The supernatant was extracted, aliquoted, and frozen at -80°C until use. Protein extracts were quantified using Pierce™ BCA Protein Assay Kit according to the manufacturer’s instructions.

*D. rerio* used for the long-term feeding scheme were kept in 3-L tanks (20-25 fish/tank). In addition to their daily standard feed including *Artemia*, fish in these tanks were also fed ∼500 transgenic *D. melanogaster* larvae (or larval extract) every week for ∼3 months. Individuals were selected at random for tissue extraction for each round of experiment of westerns, histology, and Mass Spectrometry. Experiments were repeated two to three times. Whole brain tissue was processed in the same manner as the gut tissue described above.

### 2.3. Histology

*D. melanogaster* larvae were prepared for histology by immersion in 4% PFA prepared in 1x PBS overnight at 4°C. *D. rerio* tissues were prepared for histology in the same manner, with the addition of a ventral incision. Histology work was performed at the Advanced Molecular Pathology Laboratory (AMPL @ IMCB, A*STAR) using standard protocols. Briefly, *D. rerio* tissues were decalcified in OSTEOSOFT® and trimmed to the appropriate region. After decalcification, all tissues were dehydrated in an ascending series of ethanol, cleared with xylene, and then embedded in paraffin wax. Five-micrometer sections were cut and placed onto glass slides. *D. rerio* slides were dewaxed in xylene and hydrated with a descending series of ethanol before being stained with hematoxylin and eosin. After which, the slides were dehydrated through an ascending series of ethanol to xylene before being coverslipped.

Unstained sections were stained with congo red as follows [46]. Briefly, slides were dewaxed in xylene for one hour, hydrated with a descending series of ethanol, stained for 20 minutes in filtered 0.5% w/v congo red in 50% ethanol, differentiated in 1% sodium hydroxide in 50% ethanol for 30 seconds, dehydrated through an ascending series of ethanol to xylene and coverslipped in DEPEX mounting medium.

Congo red fluorescence images were captured with an Axio Observer.Z1/7 LMS800 confocal microscope system under the following conditions; Excitation wavelength 488 nm and detection with a bandpass filter 600-700 nm. Images were analyzed using ZEISS ZEN software.

### 2.4. Sample preparation for western blotting

Protein samples from zebrafish gut and brain tissue extracts were normalized to equal concentrations and used for SDS-PAGE. As trial experiments showed that total protein can degrade during gut tissue extraction despite protease inhibitors, experiments were conducted in biological triplicates, and shown in duplicates in figures. Protein from ten *D. melanogaster* larvae per condition was used for *Drosophila* samples. The soluble fraction of the tdTomato larvae yielded 4.8 μg/μl total protein and the Aβ larvae yielded 3.5 μg/μl when extracted in 300ul. The following quantities of protein were loaded per lane, per sample: 8 μg of fly larvae and zebrafish brain samples, and 20 μg of zebrafish gut samples. Protein samples were denatured in 1x laemmli buffer with 50mM DTT at 80°C for eight minutes and loaded onto Biorad Mini-Protean 4-15% TGX precast protein gels to run for 30 minutes at 200 V under reducing conditions. Proteins were transferred to PVDF membranes in tris-glycine plus 20% ethanol transfer buffer for 90 minutes at 90 V. Following the transfer, membranes were rinsed in 1x PBS and blocked overnight at 4°C in 4% non-fat milk in 1x PBST 0.1%. Primary antibody in 1% non-fat milk, 1x PBST 0.1% (BioLegend Anti-β-Amyloid, 1-16 Antibody, clone 6E10) was added at 1:7500 for 90 minutes at room temperature, then washed three times in 1x PBS. Membranes were then incubated in secondary antibody in 1% non-fat milk, 1x PBST 0.1%, 1:10000 (Aligent Dako Anti-mouse, HRP-conjugated) for one hour at room temperature. The secondary antibody was removed by five, ten-minute washes in 1x PBS, and then membranes were incubated in a 1:1 mix of Thermo Fisher SuperSignal™ West Atto ECL substrate for 5 minutes. The excess substrate was drained, then membranes were stored in plastic wrap and imaged.

### 2.5. Mass spectrometry

Mass spectrometry analyses were performed by the Protein and Proteomics Centre, Department of Biological Sciences, National University of Singapore. In brief, protein lysate samples were processed using the S-Trap micro column (Protifi) according to the manufacturer’s recommendations, and LCMS data was acquired on an Eksigent NanoLC-Ultra & SCIEX TripleTOF 6600. Comprehensive details of the run conditions are provided in the supplementary information (S2 Mass spec. methods). Data analysis was performed using ProteinPilot 5.03 (SCIEX) using the UniProt *Drosophila melanogaster* and *Danio rerio* reference proteome databases and spiked with common contaminant proteins (cRAP), and custom peptide search sequences for Aβ, CRY2, and mCherry (S2 Mass spec methods). The following parameters were used: thorough search mode, MMTS (lysates), trypsin enzyme, common biological modifications enabled, and detected protein threshold score of 0.05.

### 2.6. Protein multiple sequence alignment

Multiple protein sequence alignments were performed using NCBI BLASTp and visualized using NCBI Multiple Sequence Alignment Viewer 1.22.2. The following ENSEMBL database records were used for the alignment: Human ENSP00000284981, Mouse lemur ENSMICP00000039774, White-tufted-ear marmoset ENSCJAP00000067269, Mouse ENSMUSP00000154061, Zebrafish ENSDARP00000110143.

## 3. Results

To examine if transmission of Aβ by ingestion can occur, we used a *Drosophila melanogaster* transgenic line that expressed a fluorescently tagged optogenetic Aβ1-42 peptide (human Aβ-CRY2-mCh) and oligomerizes into aggregates [23,43]. A second transgenic, fluorescent line expressing TdTomato served as a control [43]. Experimental or control larvae were then administered as feed for adult zebrafish that readily consume insect larvae (S1 Feeding video). Mass spectrometry, western blotting, and histological analysis were used to evaluate Aβ transmission, retention, accumulation, and associated pathologies. Two feeding schemes were used, one intensive and short-term over 48 hours to check for survival and retention of Aβ in the digestive system, and a second less intensive and long-term over three months to assay for accumulation and uptake into the brain tissue (Figure 1).

**Figure 1.**
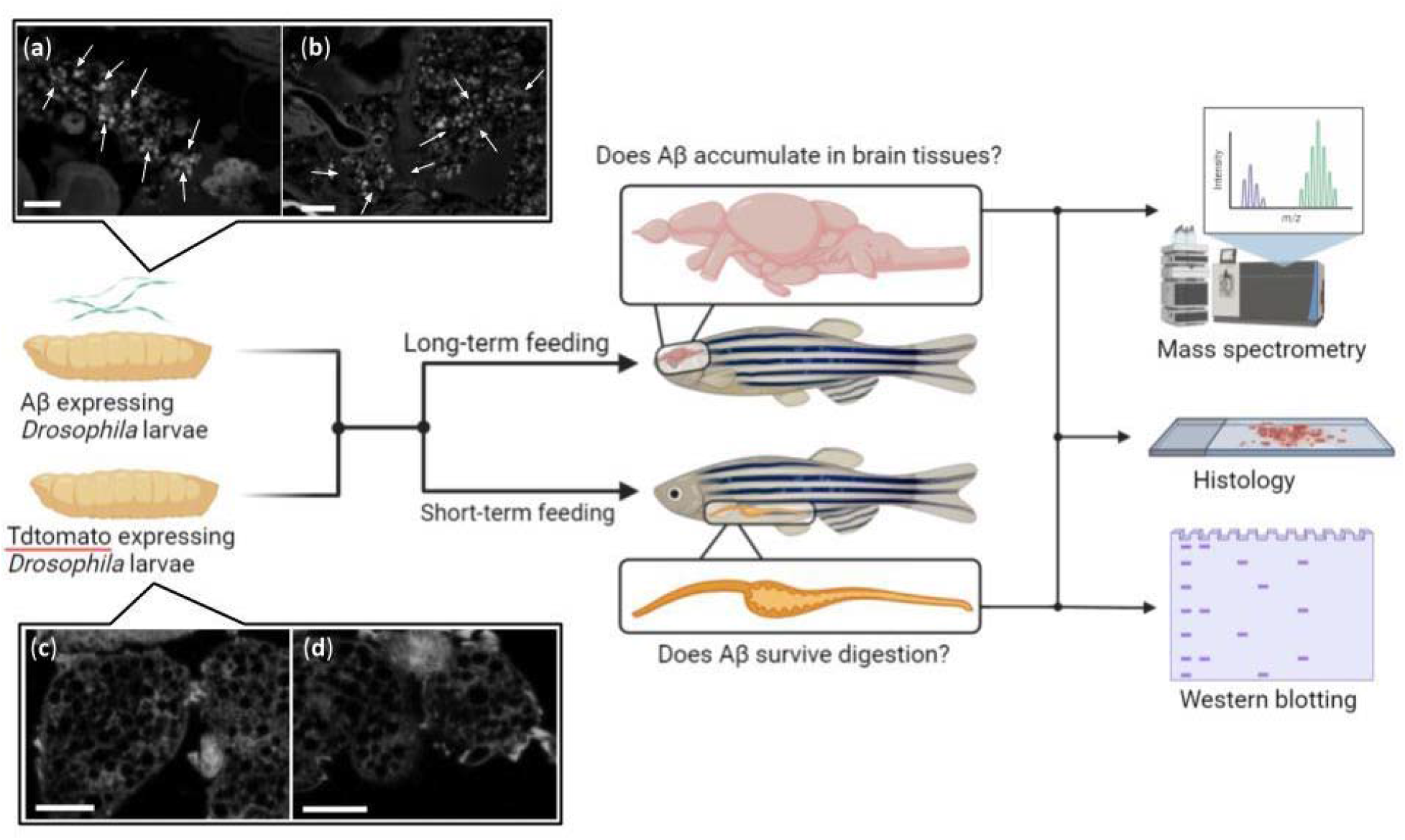
Schematic overview of the Aβ transmissibility study. (**a**) & (**b**) Congo red stained paraffin sections of transgenic *D. melanogaster* larvae expressing human Aβ fusion protein (Aβ-CRY2 - mCherry), n = 3. White arrows indicate amyloid deposits. (**c**) & (**d**) Congo red stained sections of transgenic *D. melanogaster* larvae expressing tdTomato, n = 3.

As previous studies had used live imaging to visualize Aβ aggregates in *Drosophila* larvae [23,43], we first tested if they were detectable by methods that can be used in this study such as histology and western blotting. Congo red staining on histological sections showed a distinct staining pattern in Aβ expressing *D. melanogaster* larvae (experimental larvae; Figure 1a,b) compared to transgenic, fluorescent larvae used as controls (Figure 1c,d).

Similarly, the aggregates were also detectable by western blotting at a feed-relevant scale where the protein samples prepared from 10 larvae were diluted 300 fold (Figure 2a,b). On the western blots, the Aβ was detectable either in the form of an ∼88.5 kDa or an ∼61.5 kDa band corresponding to the full Aβ-CRY2-mCh or the Aβ-CRY2 respectively (Figure 2a) but were not detectable as Aβ peptide (∼10 kDa; Figure 2b). Therefore, Aβ aggregates, when present, even as a tiny fraction of the total protein, are readily detectable by these methods (See Supplemental Material S3).

**Figure 2.**
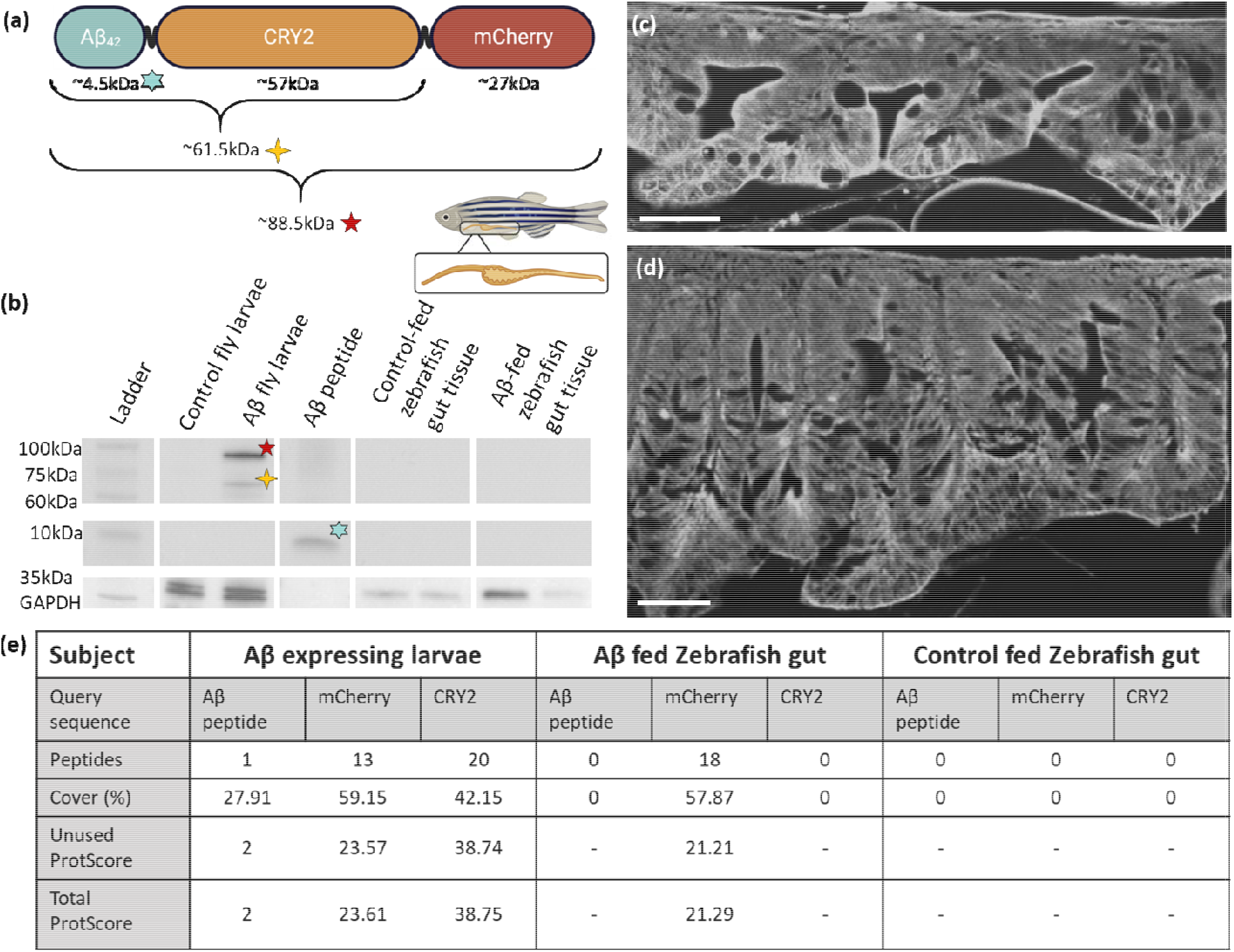
Short-term feeding scheme searching for retention of Aβ in the gut tissue of *D. rerio*. (**a**) Schematic diagram of tissue extract regions, and estimated molecular weights of individual and combined components of the Aβ fusion protein expressed by transgenic *D. melanogaster* larvae. (**b**) Aβ (clone 6e10 antibody) western blotting of *D. rerio* gut tissue 24 hours after short-term feeding, 20 μg, n = 3 per condition. Aβ larvae = *D. melanogaster* larvae expressing Aβ fusion protein, 8 μg, n = 2. tdTomato larvae = *D. melanogaster* larvae expressing tdTomato, 8 μg, n = 2. Aβ peptide **=** human amyloid beta 1-42 peptide, 0.5 μg, n = 2. (c & d) Representative congo-red amyloid stained paraffin sections of (**c**) *D. rerio* gut 24 hours after feeding with larvae expressing tdTomato, n = 3, and (**d**) *D. rerio* gut 24 hours afte**r** feeding with larvae expressing Aβ42 n = 3. Scale bars = 50 μm. (e) Detection of target proteins via MS in assorted tissues. Representative samples with the highest ProtScore, n = 3 per condition. ProtScore >1.64 = < 1 % Local false discovery rate. ProtScore >0.47 = <1 % Global false discovery rate. ‘-’ indicates no peptide evidence detected fo**r** analysis. Peptide confidence >95%. Analysis performed in ProteinPilot software 5.03.

For transmissibility via ingestion, Aβ would have to evade the digestive processes following intake. If it does so, it should be retained in the gut tissue lining the digestive tract after ingestion. We appraised if such retention happened by examining the gut tissue of the short-term fed zebrafish. In this scheme, approximately 20 larvae were fed to each fish over 48 hours, as described in the methods. We estimated that each larva contained ∼3 μg Aβ-CRY2-mCh (Table S3.1). If the survival and retention rates of Aβ in the gut corresponded to the behavior of other prions such as PrP^sc^ where retention after oral administration has been quantified [47], we would expect ∼3.9 μg of the fusion protein to remain in the whole gut tissue 24 hours after feeding on 20 larvae (see Table S3.2 for details). This would result in ∼75 ng Aβ containing protein in the gut samples reported in Figure 2b (also see Table S3.3). This amount is well within the detection threshold of our western blots (Figure S3.2). However, Aβ was undetectable in the gut tissue of these fish by western blotting at any of the molecular weights (∼88.5 kDa, ∼61.5 kDa, ∼10 kDa; Figure 2b), or by Congo red staining in histological sections (Figure 2c,d). To further confirm these observations we also performed LC-MS of the gut samples. LS-MS detected the mCherry portion of the Aβ fusion protein in two out of three samples, but not the Aβ or CRY2 portions (Figure 2e). Given the variability between individual samples, it would seem that mCherry residue detection may be a consequence of incomplete digestion as opposed to retention (Table S2.3). Thus, although amyloid and Aβ-specific staining was present in the food (*D. melanogaster* larvae), it was not present in a detectable quantity in the gut tissue of the zebrafish eating the larvae.

While no gut retention of Aβ was observable in the short-term feeding period, the possibility remained for it being present at a sub-detection threshold level initially, which eventually accumulates and leads to amyloidosis in the brain in the longer term. Further, it is also possible that the western blotting and histological techniques are not sensitive enough to detect such an accumulation. To test thes possibilities, we employed a long-term feeding scheme and added a more sensitive Mass Spectrometry method in our next experiment. For this purpose, adult fish tanks (20-25 fish/tank) were also fed ∼50 transgenic *D. melanogaster* larvae, or larval extracts a week (control and experimental) for ∼ 3 months. No gross changes in the brain, or common indicators of Aβ pathologies seen in other animals, such as plaques, lesions, and hemorrhages [48,49] were observed in these fish regardless of the feed being control or experimental larvae (Figure 3a). Absences of Aβ aggregate staining by Congo red in histological sections of the brain tissue reinforced this observation (Figure 3b,c). Finally, Aβ was not observed in the brain tissues of these animals in western blotting (Figure 3d) or Mass Spectrometry (Figure 3e) although both techniques were effective in detecting the same in the feed. Thus, no pathological seeding in a detectable form occurred in this period after repeated consumption of contaminated feed containing Aβ aggregates.

**Figure 3.**
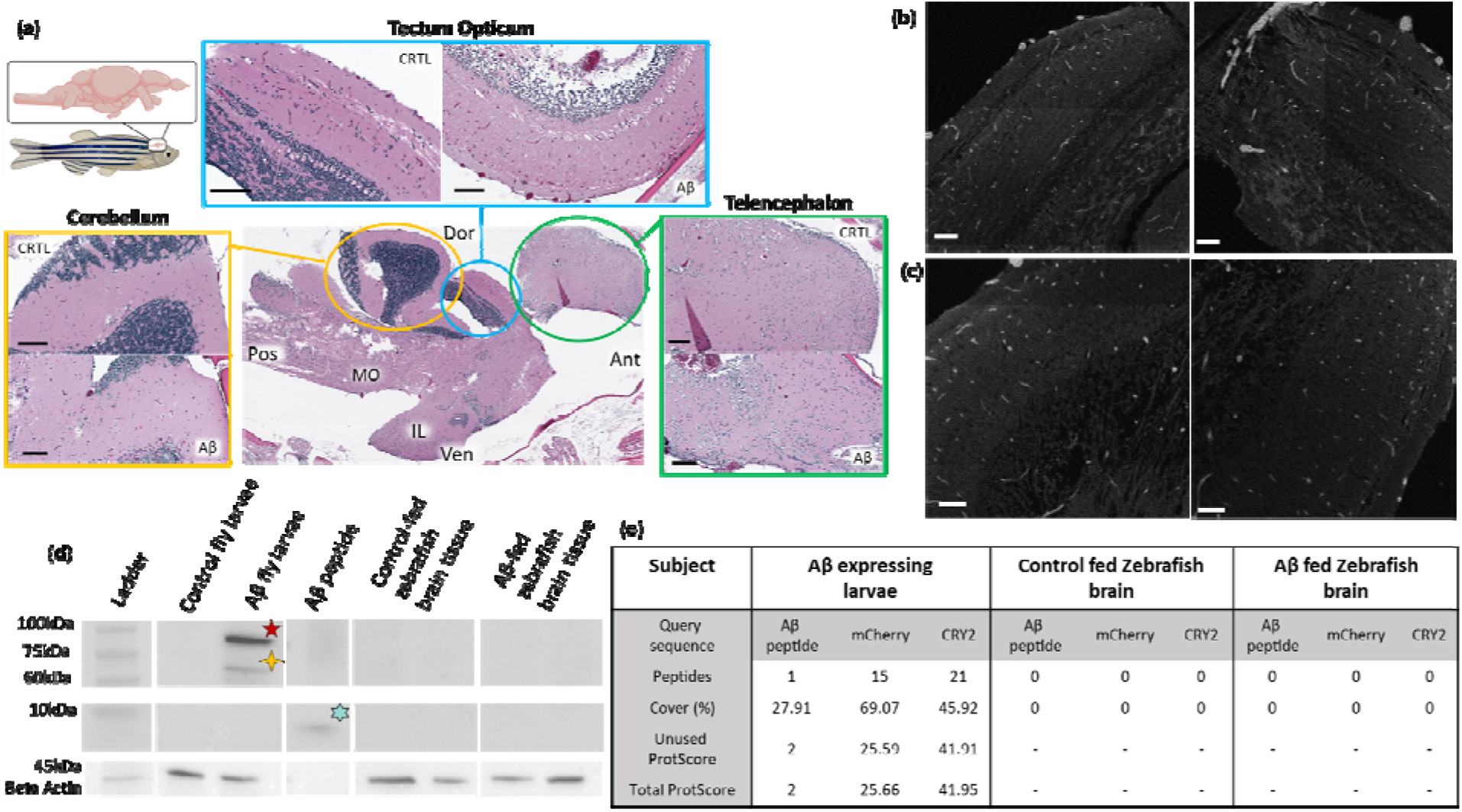
Long-term feeding scheme searching for accumulation of Aβ in the brain tissue of *D. rerio*. (**a**) Representative hematoxylin and eosin stained sections of *D. rerio* brain tissues after feeding for three months with transgenic *D. melanogaster* larvae. Aβ: Tissue of *D. rerio* fed larvae expressing Aβ; CTRL: Tissue of *D. rerio* fed larvae expressing tdTomato, MO: medulla oblongata, IL: inferior lobe, Dor: dorsal, Ven: ventral, Ant: anterior, Pos; posterior. n = 2 per condition Scale bars 100 μm. (**b & c**) Representative congo-red amyloid stained paraffin section**s** of (**b**) *D. rerio* optic tectum tissue after three months of feeding with larvae expressing tdTomato, n = 2, and (**c**) larvae expressing Aβ42 n = 2. Scale bars = 50 μm. (d) Aβ (clone 6e10 antibody) western blotting of *D. rerio* brain tissue afte**r** long-term feeding for three months, n = 2 per condition, 8 μg. Aβ larvae = *D. melanogaster* larvae expressing Aβ fusion protein, n = 2, 8 μg. tdTomato larvae = *D. melanogaster* larvae expressing tdTomato, n = 2, 8 μg. Aβ peptide = human amyloid beta 1-42 peptide, 0.5 μg, n = 2. (**e**) Detection of target proteins via MS in assorted tissues. Representative samples with the highest ProtScore n = 2 (pooled) per condition. ProtScore >1.64 = < 1 % Local false discovery rate. ProtScore >0.47 = <1 % Global false discovery rate. ‘-’ indicates no peptide evidence detected for analysis. Peptid confidence >95%. Analysis performed in ProteinPilot software 5.03.

## 4. Discussion

Prion-mediated diseases are well-reported as zoonotically transmissible via the digestive tract [35–38]. The retention of Aβ in the gut tissue represents a potential route for pathogenicity and migration to the brain, as recently suggested for both Aβ and the aggregation-prone alpha-synuclein [38,42]. The critical question is the “survival” rate or the retention of Aβ following infrequent ingestion. In the short-term feeding scheme, a 24-hour gap was left between the final feeding and extraction of gut tissue for analysis. 24 hours of fasting is expected to clear almost all of the gut content of any undigested remains in larvae as well as adult zebrafish [50–52]. Thus, it was expected that any Aβ staining seen after 24 hours is likely to come from the fish gut tissue. However, no staining was seen in either the histological sections showing the gut or western blots of the entire gut tissue. Only the Mass Spectrometry showed remnants post digestion as the mCherry component of the fusion protein Aβ-CRY2-mCherry could be detected in 2 out of 3 samples. It seems likely that most of the *D. melanogaster* larvae were digested and/or excreted to below detectable quantities, or were sufficiently degraded beyond the point of specific detection in the zebrafish tissue (Figure 2).

It is already promising that the surrounding fish digestive tract lining tissue cells were free of Aβ aggregates. However, some strains of Aβ peptides derived from AD brains are more vulnerable to denaturation and degradation by proteases than others [17]. A more resistant form of Aβ peptide could potentially survive the digestive process long enough to transmit to the gut tissues at a sub-detection threshold, only to become a pathogenic seed once it migrates to the brain [42]. The long-term feeding assay where fish were routinely fed the same transgenic *D. melanogaster* larvae over three months can address these possibilities. Synthetic human Aβ is reported to aggregate, and cause noticeable brain and behavioral impairments when inoculated intracerebrally into larval zebrafish [31,33,46,53]. While the Aβ fusion protein expressed in larvae used as feed in this study (Aβ-CRY2-mCherry) is much larger and quite different in comparison, we previously demonstrated that even this form of Aβ exhibits many of the same pathological properties. It reduced the lifespan of *D. melanogaster* and *C. elegans* and caused physical neuronal damage in *D. rerio* [23]. The pathologies, and irreversible nature of these aggregates compared with CRY2 alone or other controls, indicated that the fusion protein is a reasonable approximation of damage due to Aβ aggregation [23].

The brain tissue of the long-term-fed fish, however, showed no indication of Aβ presence or associated pathologies (Figure 3). Our choice of 3 months of feeding was based on the Aβ inoculation studies, where zebrafish tend to display symptoms and detectable changes within weeks of exposure. The synthetic Aβ load in these cases is likely to be many-fold higher than the hypothetical scenario of ingested undetectable seeds [31,33,46,53] that the design of our study intended to mimic as “natural” conditions. Hence, it is reassuring that no signs of Aβ aggregation or its prion-like features occurred at this time.

The potential for undetectable seeds acting as nucleation sites with pathologies developing only much later in life (years) however, remains in the experiment above. In rodents, symptoms can take upwards of four months to manifest, even in genetically predisposed transgenic lines, while primates only develop signs of cerebral amyloidosis one year or more following inoculation with AD-afflicted brain tissue [21,22,24,26,28–31]. As much as this could be a factor of the longevity of a species, transmissibility of Aβ via ingestion in *D. rerio* here also, if at all occurring, may parallel these mammalian studies where a small seeding event takes long incubation times (years). If this is the case, longer observation periods over the lifespan of the fish (∼3 years) will be necessary to completely rule out this possibility.

Another pertinent question at this stage is whether human Aβ peptide is seed-competent in triggering a misfolding cascade interspecifically in *D. rerio*. While almost all the historical research into the transmissibility of Aβ is anthroponotic, the principle of cross-species pathogenicity is highly relevant, especially in the context of protein self-propagation. The degree to which a prion from a donor species is robust to interspecific amino acid sequence differences may influence its ability to trigger a cascade in the host. In other prion diseases such as BSE or chronic wasting disease, where zoonotic transmission is documented, interspecific prion protein sequence conservation is ∼90% [35–38,54,55]. In primate Aβ studies, the long incubation times and gradual spread through the wild-type brain indicate that the Aβ peptides are likely acting as seeds and triggering misfolding of host native peptides [21,22,24,26–28]. However, the amino acid sequence identity of the Aβ region of APP in these species is 100% (Figure 4), thus, a species barrier does not pose a factor in preventing this spread.

**Figure 4.**
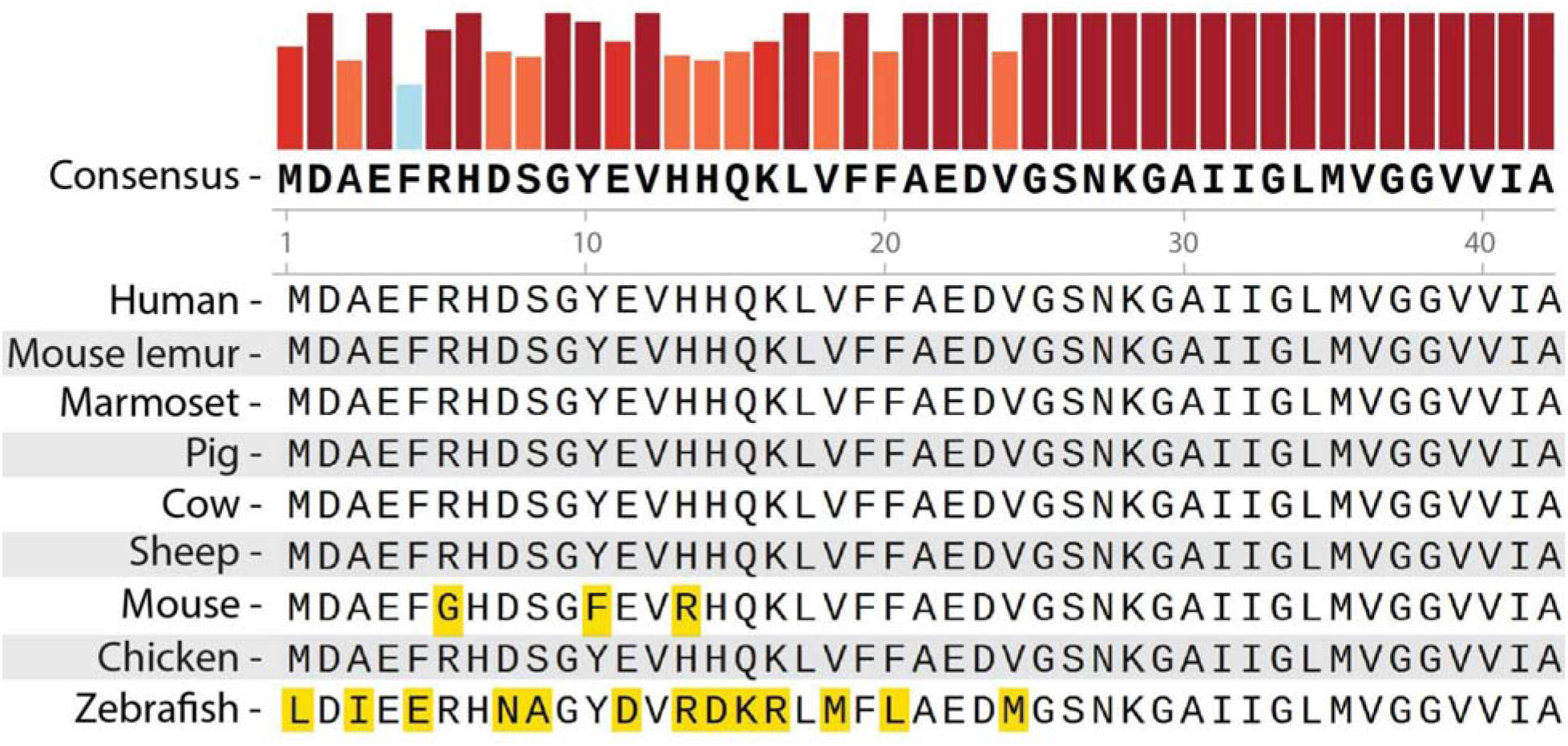
Multiple sequence alignment of amyloid precursor protein orthologs from species used in Alzheimer’s disease studies [17,21–26,28,31], and common livestock animals, focused on the human Aβ peptide region. Human: *Homo sapiens*, Marmoset: *Callithrix jacchus*, Mouse lemur: *Microcebus murinus*, Pig: *Sus scrofa*, Cow: *Bos taurus*, Sheep: *Ovis aries*, Mouse: *Mus musculus*,Chicken: *Gallus gallus*, Zebrafish: *Danio rerio*. Generated using Snapgene 7.1. Bars above the consensus sequence represent the consensus percentage (0-100%). An orange square highlights an amino acid variance from the human peptide sequence.

Mouse models, on the other hand, do not display Aβ pathologies post-inoculation with human AD-afflicted brain tissue unless they are also altered transgenically to express human APP. Knock-out of the native mouse APP in these transgenics increases the deposition rate of Aβ [21,25]. The Aβ region differs by only three amino acids in mice and highlights the need for high sequence similarity for Aβ to act as a prion (Figure 4). Fish APP orthologues are only 63-70% similar to human APP_695_ and are especially variable in the region forming the Aβ peptide (Figure 4) [56]. Hence, the absence of Aβ seeding in our experiments may be simply a reflection of this sequence dissimilarity rather than the inability of Aβ to act as a prion seed via ingestion. Furthermore, there is a paucity of information regarding the native aggregation capacity of endogenous zebrafish Aβ, with no evidence indicating it is capable of misfolding or forming plaques akin to human Aβ peptides. We chose to conduct the experiments as reported because unlike most mammalian models tested, the synthetic Aβ could simulate pathologies relevant to AD in *D. rerio* [31,46,53]. However, it was difficult to know whether or not the synthetic Aβ also misfolded native *D. rerio* Aβ peptide. Our results suggest that host Aβ misfolding in fish from human Aβ aggregates is unlikely. This is encouraging, as along with previous rodent studies [21,25], our study suggests that transmission via ingestion across species may not be possible unless Aβ sequences are identical.

At present, we can only speculate about the consequences of brain extracts from AD patients being consumed directly, or Aβ plaques forming in farm animals used as food. An additional factor to note when considering the potential of transmission by ingestion in other species is variation in digestive efficiency, as well as the “leakiness” of the gut to blood to the brain. Zebrafish show functional conservation of the blood-brain barrier (BBB) with mammals [57] and the average digestion efficiencies of fish and non-ruminant mammals are reportedly similar when consuming whole invertebrates [58]. Hence, these may not be a significant factor extrapolating the outcomes reported here. While the consumption of human brain tissue containing Aβ is a highly unlikely scenario, the strong conservation of the Aβ peptide sequence between livestock animals and humans may warrant further investigation into this potential mode of transmission (Figure 4.).

## 5. Conclusions

In conclusion, it appears that while iatrogenic transmission of AD and other neurodegenerative disorders is a serious health risk, inter-species ingestion of Aβ may be a less likely route of transmission especially when Aβ sequence similarity between species is low. If gut retention of Aβ and transmission to the brain was occurring at all, it fell below detectable levels. Further experimentation will be required though to determine if Aβ pathology arises over a longer time scale of years, and whether pathogenicity of human, or livestock animal, Aβ via the human digestive tract is possible.

## 6. List of abbreviations

Aβ: amyloid beta peptide
Aβ_42_: Human amyloid beta peptide 1-42
AD: Alzheimer’s disease
APP: amyloid protein precursor

## 7. Additional Files

‘S1 - Feeding video’, video file (.mp4). Zebrafish feeding. Contains video footage of zebrafish consuming both control and Aβ-expressing *Drosophila* larvae.

‘S2 - Mass spec methods’, PDF document (.pdf). Full mass spectrometry methods. Contains detailed descriptions of the mass spectrometry methods, false discovery rate graphs for analysis, and peptide sequences used to query the database.

‘S3 - Detection of amyloid precursor protein and Aβ_42_’, PDF document (.pdf). Contains additional figures highlighting that native zebrafish amyloid precursor protein was detectable by western blot and mass spectrometry in the brain tissue samples, and the detection limit of Aβ_42_ peptide by western blotting. Also contains calculations estimating Aβ retention in the gut tissue if it behaves as a prion protein.

## 8. Declarations

### Ethics approval and consent to participate

Zebrafish studies were performed following protocol number IACUC#: 201571 approved by the Institutional Review Board of A-STAR laboratories Biological Resource Center (BRC).

### Consent for publication

Not applicable

### Availability of data and materials

The datasets used and/or analyzed during the current study to generate the figures are included within the article and its additional file(s). Additional data are available from the corresponding author upon request.

### Competing interests

The authors declare no competing interests.

### Funding

This research was supported by Yale-NUS College via the grant from the Ministry of Health’s National Medical Research Council’s Healthy Longevity Catalyst Award (HLCA), grant number MOH-000517-00 to A.S.M.

### Author Contributions

Conceptualization, A.S.M, and J.G; methodology, A.S.M., and J.R.; formal analysis, J.R., and A.S.M.; investigation, J.R., and A.S.M; resources, J.G., N.T., and A.S.M; writing— original draft preparation, J.R, and A.S.M.; writing—review and editing, A.S.M and J.R.; visualization, J.R.; supervision, A.S.M..; project administration, A.S.M; funding acquisition, A.S.M.

## Acknowledgments

We would like to thank the zebrafish staff at A-STAR, IMCB, and Yale-NUS College for their support. Additionally, many thanks to Dr. Ce-Belle Chen for her advice on western blotting for amyloids, and Dr. Caroline Kibat and Tanisha Goel for their assistance in sample collection/preparation.

## Notes

### Competing Interest Statement

The authors have declared no competing interest.

### Summary of Updates

Revised Figure 1; New supplementary data (S3) to support detection sensitivity and associated calculations; Revised Figure 4; Revised Discussion; Author affiliations updated; Author name spelling corrected

## References

1. LaFerla FM, Green KN, Oddo S. Intracellular amyloid-beta in Alzheimer’s disease. Nat Rev Neurosci. 2007;8:499–509.

2. Masters CL, Simms G, Weinman NA, Multhaup G, McDonald BL, Beyreuther K. Amyloid plaque core protein in Alzheimer disease and Down syndrome. Proc Natl Acad Sci U S A. 1985;82:4245–9.

3. van Dyck CH, Swanson CJ, Aisen P, Bateman RJ, Chen C, Gee M, et al. Lecanemab in Early Alzheimer’s Disease. N Engl J Med. 2023;388:9–21.

4. Weglinski C, Jeans A. Amyloid-β in Alzheimer’s disease - front and centre after all? Neuronal Signal. 2023;7:NS20220086.

5. Hawksworth J, Fernández E, Gevaert K. A new generation of AD biomarkers: 2019 to 2021. Ageing Res Rev. 2022;79:101654.

6. Kepp KP, Robakis NK, Høilund-Carlsen PF, Sensi SL, Vissel B. The amyloid cascade hypothesis: an updated critical review. Brain [Internet]. 2023; Available from: 10.1093/brain/awad159

7. Sims JR, Zimmer JA, Evans CD, Lu M, Ardayfio P, Sparks J, et al. Donanemab in Early Symptomatic Alzheimer Disease: The TRAILBLAZER-ALZ 2 Randomized Clinical Trial. JAMA [Internet]. 2023; Available from: 10.1001/jama.2023.13239

8. Chow VW, Mattson MP, Wong PC, Gleichmann M. An overview of APP processing enzymes and products. Neuromolecular Med. 2010;12:1–12.

9. Walsh DM, Selkoe DJ. A critical appraisal of the pathogenic protein spread hypothesis of neurodegeneration. Nat Rev Neurosci. 2016;17:251–60.

10. Xiao Y, Ma B, McElheny D, Parthasarathy S, Long F, Hoshi M, et al. Aβ(1-42) fibril structure illuminates self-recognition and replication of amyloid in Alzheimer’s disease. Nat Struct Mol Biol. 2015;22:499–505.

11. Törnquist M, Cukalevski R, Weininger U, Meisl G, Knowles TPJ, Leiding T, et al. Ultrastructural evidence for self-replication of Alzheimer-associated Aβ42 amyloid along the sides of fibrils. Proc Natl Acad Sci U S A. 2020;117:11265–73.

12. Wu Y-S, Huang S-J, Wu M-H, Tu L-H, Lee M-C, Chan JCC. Aβ _42_ oligomers can seed the fibrillization of Aβ _40_ peptides. J Chin Chem Soc. 2022;69:1318–25.

13. Walker LC, Schelle J, Jucker M. The Prion-Like Properties of Amyloid-β Assemblies: Implications for Alzheimer’s Disease. Cold Spring Harb Perspect Med [Internet]. 2016;6. Available from: 10.1101/cshperspect.a024398

14. Watts JC, Prusiner SB. β-Amyloid Prions and the Pathobiology of Alzheimer’s Disease [Internet]. Cold Spring Harbor Perspectives in Medicine. 2018. p. a023507. Available from: 10.1101/cshperspect.a023507

15. Condello C, Merz GE, Aoyagi A, DeGrado WF, Prusiner SB. Aβ and Tau Prions Causing Alzheimer’s Disease. Methods Mol Biol. 2023;2561:293–337.

16. Carlson GA, Prusiner SB. How an Infection of Sheep Revealed Prion Mechanisms in Alzheimer’s Disease and Other Neurodegenerative Disorders. Int J Mol Sci [Internet]. 2021;22. Available from: 10.3390/ijms22094861

17. Watts JC, Condello C, Stöhr J, Oehler A, Lee J, DeArmond SJ, et al. Serial propagation of distinct strains of Aβ prions from Alzheimer’s disease patients. Proc Natl Acad Sci U S A. 2014;111:10323–8.

18. Jaunmuktane Z, Mead S, Ellis M, Wadsworth JDF, Nicoll AJ, Kenny J, et al. Evidence for human transmission of amyloid-β pathology and cerebral amyloid angiopathy. Nature. 2015;525:247–50.

19. Purro SA, Farrow MA, Linehan J, Nazari T, Thomas DX, Chen Z, et al. Transmission of amyloid-β protein pathology from cadaveric pituitary growth hormone. Nature. 2018;564:415–9.

20. Banerjee G, Samra K, Adams ME, Jaunmuktane Z, Parry-Jones AR, Grieve J, et al. Iatrogenic cerebral amyloid angiopathy: an emerging clinical phenomenon. J Neurol Neurosurg Psychiatry [Internet]. 2022; Available from: 10.1136/jnnp-2022-328792

21. Meyer-Luehmann M, Coomaraswamy J, Bolmont T, Kaeser S, Schaefer C, Kilger E, et al. Exogenous induction of cerebral beta-amyloidogenesis is governed by agent and host. Science. 2006;313:1781–4.

22. Gary C, Lam S, Hérard A-S, Koch JE, Petit F, Gipchtein P, et al. Encephalopathy induced by Alzheimer brain inoculation in a non-human primate. Acta Neuropathol Commun. 2019;7:126.

23. Lim CH, Kaur P, Teo E, Lam VYM, Zhu F, Kibat C, et al. Application of optogenetic Amyloid-β distinguishes between metabolic and physical damages in neurodegeneration. Elife [Internet]. 2020;9. Available from: 10.7554/eLife.52589

24. Lam S, Petit F, Hérard A-S, Boluda S, Eddarkaoui S, Guillermier M, et al. Transmission of amyloid-beta and tau pathologies is associated with cognitive impairments in a primate. Acta Neuropathol Commun. 2021;9:165.

25. Steffen J, Krohn M, Schwitlick C, Brüning T, Paarmann K, Pietrzik CU, et al. Expression of endogenous mouse APP modulates β-amyloid deposition in hAPP-transgenic mice. Acta Neuropathol Commun. 2017;5:49.

26. Baker HF, Ridley RM, Duchen LW, Crow TJ, Bruton CJ. Evidence for the experimental transmission of cerebral beta-amyloidosis to primates. Int J Exp Pathol. 1993;74:441–54.

27. Maclean CJ, Baker HF, Ridley RM, Mori H. Naturally occurring and experimentally induced beta-amyloid deposits in the brains of marmosets (Callithrix jacchus). J Neural Transm. 2000;107:799–814.

28. Ridley RM, Baker HF, Windle CP, Cummings RM. Very long term studies of the seeding of beta-amyloidosis in primates. J Neural Transm. 2006;113:1243–51.

29. Bornemann KD, Staufenbiel M. Transgenic mouse models of Alzheimer’s disease. Ann N Y Acad Sci. 2000;908:260–6.

30. Sturchler-Pierrat C, Staufenbiel M. Pathogenic mechanisms of Alzheimer’s disease analyzed in the APP23 transgenic mouse model. Ann N Y Acad Sci. 2000;920:134–9.

31. Nery LR, Eltz NS, Hackman C, Fonseca R, Altenhofen S, Guerra HN, et al. Brain intraventricular injection of amyloid-β in zebrafish embryo impairs cognition and increases tau phosphorylation, effects reversed by lithium. PLoS One. 2014;9:e105862.

32. Bhattarai P, Thomas AK, Zhang Y, Kizil C. The effects of aging on Amyloid-β42-induced neurodegeneration and regeneration in adult zebrafish brain. Neurogenesis (Austin). 2017;4:e1322666.

33. Saleem S, Kannan RR. Zebrafish: an emerging real-time model system to study Alzheimer’s disease and neurospecific drug discovery [Internet]. Cell Death Discovery. 2018. Available from: 10.1038/s41420-018-0109-7

34. Javed I, Peng G, Xing Y, Yu T, Zhao M, Kakinen A, et al. Inhibition of amyloid beta toxicity in zebrafish with a chaperone-gold nanoparticle dual strategy. Nat Commun. 2019;10:3780.

35. Collinge J. Human prion diseases and bovine spongiform encephalopathy (BSE) [Internet]. Human Molecular Genetics. 1997. p. 1699–705. Available from: 10.1093/hmg/6.10.1699

36. Kong Q, Zheng M, Casalone C, Qing L, Huang S, Chakraborty B, et al. Evaluation of the human transmission risk of an atypical bovine spongiform encephalopathy prion strain. J Virol. 2008;82:3697–701.

37. Lee J, Kim SY, Hwang KJ, Ju YR, Woo H-J. Prion Diseases as Transmissible Zoonotic Diseases [Internet]. Osong Public Health and Research Perspectives. 2013. p. 57–66. Available 10.1016/j.phrp.2012.12.008

38. Liu W, Lim K-L, Tan E-K. Intestine-derived α-synuclein initiates and aggravates pathogenesis of Parkinson’s disease in Drosophila. Transl Neurodegener. 2022;11:44.

39. Brown P, McShane LM, Zanusso G, Detwile L. On the question of sporadic or atypical bovine spongiform encephalopathy and Creutzfeldt-Jakob disease. Emerg Infect Dis. 2006;12:1816–21.

40. Seitz R, von Auer F, Blümel J, Burger R, Buschmann A, Dietz K, et al. Impact of vCJD on blood supply [Internet]. Biologicals. 2007. p. 79–97. Available from: 10.1016/j.biologicals.2007.01.002

41. Cassard H, Torres J-M, Lacroux C, Douet J-Y, Benestad SL, Lantier F, et al. Evidence for zoonotic potential of ovine scrapie prions. Nat Commun. 2014;5:5821.

42. Jin J, Xu Z, Zhang L, Zhang C, Zhao X, Mao Y, et al. Gut-derived β-amyloid: Likely a centerpiece of the gut-brain axis contributing to Alzheimer’s pathogenesis. Gut Microbes. 2023;15:2167172.

43. Kaur P, Kibat C, Teo E, Gruber J, Mathuru A, Tolwinski ANS. Use of Optogenetic Amyloid-β to Monitor Protein Aggregation in and. Bio Protoc. 2020;10:e3856.

44. Nathan FM, Kibat C, Goel T, Stewart J, Claridge-Chang A, Mathuru AS. Contingent stimulus delivery assay for zebrafish reveals a role for CCSER1 in alcohol preference. Addict Biol. 2022;27:e13126.

45. Gupta T, Mullins MC. Dissection of Organs from the Adult Zebrafish [Internet]. Journal of Visualized Experiments. 2010. Available from: 10.3791/1717

46. Javed I, Peng G, Xing Y, Yu T, Zhao M, Kakinen A, et al. Inhibition of amyloid beta toxicity in zebrafish with a chaperone-gold nanoparticle dual strategy. Nat Commun. 2019;10:3780.

47. Urayama A, Concha-Marambio L, Khan U, Bravo-Alegria J, Kharat V, Soto C. Prions efficiently cross the intestinal barrier after oral administration: Study of the bioavailability, and cellular and tissue distribution in vivo. Sci Rep. 2016;6:32338.

48. Eng JA, Frosch MP, Choi K, William Rebeck G, Greenberg SM. Clinical manifestations of cerebral amyloid angiopathy-related inflammation [Internet]. Annals of Neurology. 2004. p. 250–6. Available from: 10.1002/ana.10810

49. Abrahamson EE, Kofler JK, Becker CR, Price JC, Newell KL, Ghetti B, et al. 11C-PiB PET can underestimate brain amyloid-β burden when cotton wool plaques are numerous. Brain. 2022;145:2161–76.

50. Field HA, Kelley KA, Martell L, Goldstein AM, Serluca FC. Analysis of gastrointestinal physiology using a novel intestinal transit assay in zebrafish. Neurogastroenterol Motil. 2009;21:304–12.

51. Collymore C, Rasmussen S, Tolwani RJ. Gavaging adult zebrafish. J Vis Exp [Internet]. 2013; Available from: 10.3791/50691

52. Graves CL, Chen A, Kwon V, Shiau CE. Zebrafish harbor diverse intestinal macrophage populations including a subset intimately associated with enteric neural processes. iScience. 2021;24:102496.

53. Bhattarai P, Thomas AK, Zhang Y, Kizil C. The effects of aging on Amyloid-β42-induced neurodegeneration and regeneration in adult zebrafish brain. Neurogenesis (Austin). 2017;4:e1322666.

54. Li L, Coulthart MB, Balachandran A, Chakrabartty A, Cashman NR. Species barriers for chronic wasting disease by in vitro conversion of prion protein. Biochem Biophys Res Commun. 2007;364:796–800.

55. Wopfner F, Weidenhöfer G, Schneider R, von Brunn A, Gilch S, Schwarz TF, et al. Analysis of 27 mammalian and 9 avian PrPs reveals high conservation of flexible regions of the prion protein. J Mol Biol. 1999;289:1163–78.

56. Musa A, Lehrach H, Russo VA. Distinct expression patterns of two zebrafish homologues of the human APP gene during embryonic development. Dev Genes Evol. 2001;211:563–7.

57. O’Brown NM, Pfau SJ, Gu C. Bridging barriers: a comparative look at the blood-brain barrier across organisms. Genes Dev. 2018;32:466–78.

58. Karasov WH, Douglas AE. Comparative digestive physiology. Compr Physiol. 2013;3:741–83.

