## Supplemental Methods 2 for "Evaluating the transmission risk of amyloid beta peptide via ingestion"

**Supplementary information for Raine *et al.* "Evaluating the transmission risk of amyloid beta peptide via ingestion"**

### S2 - Full mass spectrometry methods

LCMS Data Acquisition was performed under the following parameters. Eksigent NanoLC-Ultra with trap column; Trajan ProteoCol C18P 3 μm 120 Å, 300 μm x 10 mm and analytical column; Thermo Scientific Acclaim PepMap100 C18 3 μm 100 Å, 75 μm x 250 mm. The following solvent ratios were used: Solvent A: 2% acetonitrile, 0.1% formic acid and Solvent B: 98% acetonitrile, 0.1% formic acid.

Table S2.1. LC solvent scheme.

| Time | Flow rate (nL/min) | % A | % B |
|---|---|---|---|
| 0 | 300 | 95 | 5 |
| 20 | 300 | 88 | 12 |
| 60 | 300 | 70 | 30 |
| 62 | 300 | 10 | 90 |
| 69 | 300 | 10 | 90 |
| 72 | 300 | 95 | 5 |
| 85 | 300 | 95 | 5 |

SCIEX TripleTOF 5600 with the ionization parameters: Nebulizer gas (GS1) 12 units, Curtain gas 30 units, IonSpray voltage floating 2300 V, and Interface heater temperature 150 °C. MS – IDA (information-dependent acquisition) parameters: Mass range 350 – 1250 m/z, accumulation time 250 msec, maximum 20 precursors selected for fragmentation, charge state 2-4, intensity > 125 cps, dynamic exclusion for 15 sec. MSMS parameters: mass range 100 – 1800 m/z, accumulation time: 100 msec, mode High sensitivity, rolling collision energy enabled.

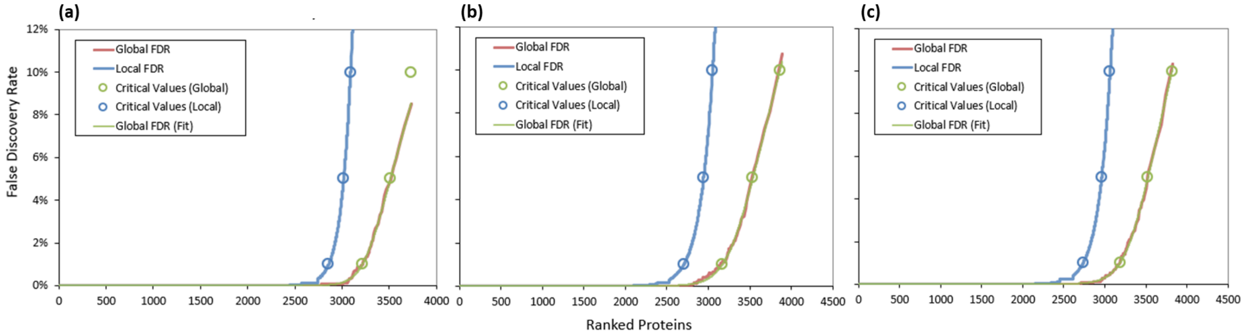


Figure S2.1. Protein false discovery rates for (**a**) Aβ expressing *D. melanogaster* larvae, (**b**) Aβ fed *D. rerio* brain tissue and (**c**) tdTomato fed *D.rerio* brain tissue. Analysis performed in ProteinPilot software 5.03.

Table S2.2. Query sequences used to search the peptide MS data. The regions highlighted indicate peptide coverage. **Green:** High confidence > 95%, **Orange:** Medium confidence 95% - 20%, **Red:** Low confidence, < 20%. Analysis performed in ProteinPilot software 5.03.

| Query label | Query amino acid sequence and peptide coverage |
|---|---|
| $A\beta_{42}$ peptide | MDAEFRHDSGYEVHHQKLVFFAEDVGSNKGAIIGLMVGGVVIA |
| mCherry | MVSKGEEDNMAIIKEFMRFKVHMEGSVNGHEFEIEGEGEGRPYEGTQTAKLKVTKG<br>GPLPFAWDILSPQFMYGSKAYVKHPADIPDYLKLSFPEGFKWERVMNFEDGGVVTVT<br>QDSSLQDGEFIYKVKLRGTNFPSDGPVMQKKTMGWEASSERMYPEDGALKGEIKQR<br>LKLKDGGHYDAEVKTTYKAKKPVQLPGAYNVNIKLDITSHNEDYTIVEQYERAEGR<br>HSTGGMDELYK |
| CRY2 | MKMDKKTIVWFRRDLRIEDNPALAAAAHEGSVFPVFIWCPEEEGQFYPGRASRWW<br>MKQSLAHLSQSLKALGSDLTLIKTHNTISAILDCIRVTGATKVVFNHLYDPVSLVRDH<br>TVKEKLVERGISVQSYNGDLLYEPWEIYCEKGKPFTSFNSYWKKCLDMSIESVMLPPP<br>WRLMPITAAAEAIWACSIEELGLENEAEKPSNALLTRAWSPGWSNADKLLNEFIEKQ<br>LIDYAKNSKKVVGNSTSLLSPYLHFGEISVRHVFQCARMKQIIWARDKNSEGEESADL<br>FLRGIGLREYSRYICFNFPFTHEQSLLSHLRFFPWDADVDKFKAWRQGRTGYPLVDAG<br>MRELWATGWMHNRIRVIVSSFAVKFLLLPWKWGMKYFWDTLLDADLECDILGWQY<br>ISGSIPDGHELDRLDNPALQGAKYDPEGEYIRQWLPELARLPTEWIHHPWDAPLTVL<br>KASGVELGTNYAKPIVDIDTARELLAKAISRTREAQIMIGAAARDPP |

Table S2.3. Full detection of target proteins via MS in assorted tissues from figure 2. ProtScore >1.64 = < 1 % Local false discovery rate. ProtScore >0.47 = <1 % Global false discovery rate. '-' indicates no peptide evidence detected for analysis. Peptide confidence >95%. Analysis performed in ProteinPilot software 5.03.

| Subject | Aβ fed Zebrafish gut #1 | | | Aβ fed Zebrafish gut #2 | | | Aβ fed Zebrafish gut #3 | | | Control fed Zebrafish gut #1, #2, #3 | | |
|---|---|---|---|---|---|---|---|---|---|---|---|---|
| Query sequence | Aβ peptide | mCherry | CRY2 | Aβ peptide | mCherry | CRY2 | Aβ peptide | mCherry | CRY2 | Aβ peptide | mCherry | CRY2 |
| Peptides | 0 | 0 | 0 | 0 | 18 | 0 | 0 | 10 | 0 | 0 | 0 | 0 |
| Cover (%) | 0 | 0 | 0 | 0 | 57.87 | 0 | 0 | 26.81 | 0 | 0 | 0 | 0 |
| Unused ProtScore | - | - | - | - | 21.21 | - | - | 8.25 | - | - | - | - |
| Total ProtScore | - | - | - | - | 21.29 | - | - | 8.28 | - | - | - | - |

| Subject | Aβ expressing larvae #1 | | | Aβ expressing larvae #2 | | | Control expressing larvae #1 | | | Control expressing larvae #2 | | |
|---|---|---|---|---|---|---|---|---|---|---|---|---|
| Query sequence | Aβ peptide | mCherry | CRY2 | Aβ peptide | mCherry | CRY2 | Aβ peptide | mCherry | CRY2 | Aβ peptide | mCherry | CRY2 |
| Peptides | 1 | 13 | 20 | 1 | 13 | 20 | 0 | 7 | 0 | 0 | 6 | 0 |
| Cover (%) | 27.91 | 59.15 | 42.15 | 27.91 | 58.72 | 44.14 | 0 | 26.66 | 0 | 0 | 27.23 | 0 |
| Unused ProtScore | 2 | 23.57 | 38.74 | 2 | 23.64 | 37.98 | - | 4.04 | - | - | 6.02 | - |
| Total ProtScore | 2 | 23.61 | 38.75 | 2 | 23.66 | 38.01 | - | 4.04 | - | - | 6.03 | - |
