## Supplemental Information 3 for "Evaluating the transmission risk of amyloid beta peptide via ingestion"

**Supplementary information for Raine *et al.* "Evaluating the transmission risk of amyloid beta peptide via ingestion"**

*Detection of amyloid precursor protein and A$\beta_{42}$*

**Figure S3.1** Shows that both methods used in this study, Western blotting and Mass Spectrometry can detect zebrafish native amyloid precursor protein B (Z-appB) in general.

(a)

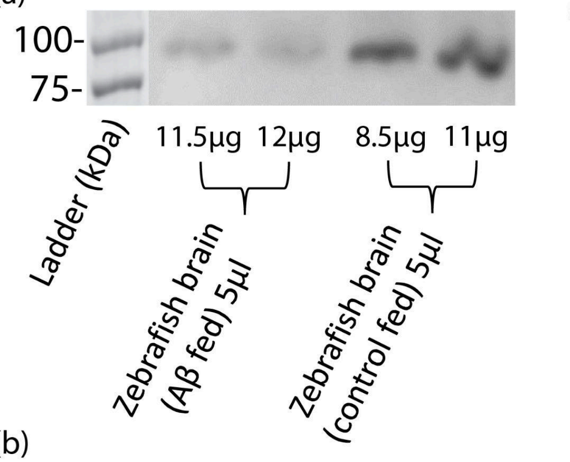


(b)

| Subject | Control fed Zebrafish Brain |
|---|---|
| Query sequence | Danio rerio amyloid precursor protein B |
| Peptides | 1 |
| Cover (%) | 2.3 |
| Unused ProtScore | 2.26 |
| Total ProtScore | 2.3 |

**Figure S3.1. Detection of native zebrafish app. (a)** Western blot with Invitrogen, beta Amyloid Polyclonal Antibody CT695 detects native Z-app B in the zebrafish brain (in the same tissue samples reported in Figure 3). **(b)** Table shows Z-appB peptides detected in Mass Spectrometric analyses of zebrafish brain tissue. ProtScore > 1.64 = < 1 % Local false discovery rate. ProtScore > 0.47 = <1 % Global false discovery rate. Peptide confidence > 95%. Analysis performed in ProteinPilot software 5.03.

**Figure S3.2** shows that ~50-100 ng of synthetic A$\beta_{42}$ peptide can be detected by the Western blotting techniques used in this study.

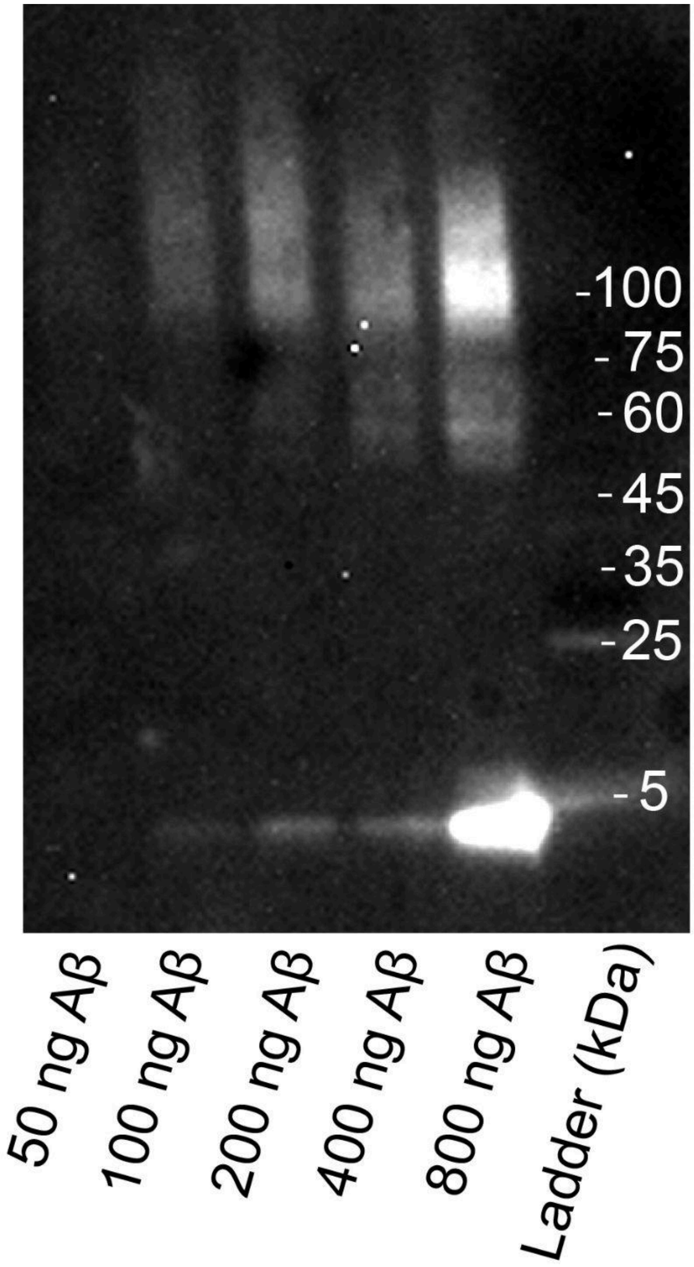


**Figure S3.2. Detection threshold of A$\beta_{42}$.** Gel electrophoresis and blotting methodology as described in Methods section 2.4. A$\beta_{42}$ peptide can be observed at 4 kDa, while aggregates that remain after the reduction and denaturation process appear on the blot as a smear (45 kDa or higher).

*Estimation of the amount of Aβ expected to be retained in the gut tissue.*

We estimated the amount of human Aβ containing transgenic protein in a *D.melanogaster* larva by quantifying the area under the peak on the Western blot and comparing it with the area under the peak for synthetic $A\beta_{42}$ peptide (ImageJ, gel analyzer, area under the peak). This analysis suggests that a single larva contains ~225ng of Aβ peptide in the 8μg total protein loaded. Thus, it accounts for ~3.5% of the total protein loaded. Assuming this ratio is representative of the total larval extracted protein (3.5 μg/μl, 300 μl), each larva would contain an average of ~3 μg of Aβ.

**Table S3.1. Estimation of average Aβ quantity per *Drosophila* larva based on quantification from Western blots.**

| Aβ per larva per 8µg of total protein (ng) | AB per ng of total protein loaded (ng) | Average total protein in 300 µl extract of 10 larvae (ng) | Average total protein per larva (ng) | Average Aβ per larva (ng) |
|---|---|---|---|---|
| 224.03 | 0.028 | 1050000 | 105000 | 3036.53 |

If Aβ behaves in a manner that is similar to the prion protein (PrP) as described by Urayama *et al.* in surviving the digestive process after oral administration, then we can expect ~4% in the stomach, ~2% in the intestines, and ~0.4% in the blood of the total orally administered protein to be retained [1]. Based on these assumptions, we estimated the amount of Aβ expected to be retained during the short-term feeding scheme in the guts 24-48 hours after having been fed with 20 *D. melanogaster* larvae in Table S3.

**Table S3.2. Estimation of Aβ expected to be retained in the zebrafish gut after the short-term feeding scheme.** Table follows PrP retention data from [1]

| Tissue | Aβ expected to be retained per larva consumed (ng) | Aβ expected to be retained following short-term feeding (ng) |
|---|---|---|
| Blood (0.4%) | 12.15 | 242.92 |
| Stomach (4%) | 121.46 | 2429.23 |
| Intestines (2%) | 60.73 | 1214.61 |
| Combined (6.4%) | 194.349 | 3886.76 |

In Table S3.3, we predict the amount of Aβ that would have been present in the Western blot as a proportion of the total gut protein loaded (shown in Figure 2).

**Table S3.3. Estimations of Aβ predicted to be present per lane of 20 μg zebrafish gut tissue if Aβ survived digestion to the same degree as PrP.**

| Average gut total protein extract (ng) | Aβ retained (ng) | Aβ to total protein (%) | Predicted average Aβ in lane (ng) |
|---|---|---|---|
| 992225 | 3886.76 | 0.39 | 78.34 |

Under the assumption that Aβ aggregates behaved akin to PrP and the digestive efficiencies of fish and non-ruminant mammals are similar when consuming whole invertebrates [2], we would expect 50-100 ng of Aβ to be present in the western blots in the high-intensity, short-term feeding scheme. This is well within the range of detection of both methods used in this manuscript. Thus, Figures and Tables in the S3

Supplement suggest that the methods used in this manuscript are adequately sensitive to detect Aβ peptide if it was present.

**References**

1. Urayama A, Concha-Marambio L, Khan U, Bravo-Alegria J, Kharat V, Soto C. Prions efficiently cross the intestinal barrier after oral administration: Study of the bioavailability, and cellular and tissue distribution in vivo. Sci Rep. 2016;6:32338.

2. Karasov WH, Douglas AE. Comparative digestive physiology. Compr Physiol. 2013;3:741–83.
